# Pangenome References Improve Biomarker Estimation from Tumor Sequencing Data

**DOI:** 10.1101/2024.11.14.623554

**Authors:** Elif Arslan, Deniz Turgut, Özem Kalay, Sinem Demirkaya-Budak, Gungor Budak, Amit Jain

## Abstract

It has recently been shown that patients from non-European ancestries are at a higher risk of inappropriate clinical intervention because of inaccurate biomarker estimation, arising from the reference bias inherent in standard methods for determining the tumor genome from sequencing data. Here we demonstrate that these inaccuracies can be reduced by using a pangenome reference appropriate for the patient’s population.

We constructed a novel secondary analysis workflow where the pangenome reference serves as a scaffold for mapping the sequencing reads, and is also included in the relevant ‘panel of normals’ needed to discriminate between germline and somatic mutations in tumor-only sequencing. This approach detects known somatic mutations in tumor-only sequencing more accurately than the standard GATK somatic calling workflow, prevalent in diagnostic settings for analysis of sequencing data from tumor-only assays, on a standard benchmark tumor sample, HCC1395 (33% relative increase in F1 score).

We also assessed the expected clinical impact of our approach by comparing the Tumor Mutational Burden (TMB) calculated from missense somatic mutations called in tumor/normal samples from 6 patients self-reported as belonging to African, 1 to Asian and 3 to European populations respectively. We find that the TMB values calculated from the tumor-only sequencing data analyzed by our workflow more closely approximate the TMB values calculated from the tumor-normal analysis of the same sample, being 35% higher on average, whereas GATK tumor-only analysis generates TMB values 56% higher on average than the tumor-normal analysis of the same sample. Tumor-normal TMB values calculated by the two methods do not vary as drastically, GATK generated values being 13% higher on average, indicating that GATK tumor-only analysis leads to significant overestimation of TMB values, which can be largely corrected by using our workflow when tumor-normal sequencing is not available.

These results indicate that pangenome based analysis has the potential to become the new standard for unbiased processing of somatic sequencing samples, following on from its increased adoption for germline sequencing analysis.

---

A recent study highlights significant disparities in tumor mutational burden (TMB) estimation and immune checkpoint inhibitor (ICI) treatment outcomes across different ancestries (Nassar et al., 2022). Using tumor-only genetic sequencing data, the researchers found that TMB estimates are inflated in non-European populations due to inadequate representation of these groups in germline reference databases. This overestimation particularly affected patients of Asian and African descent, often leading to their misclassification as TMB-high and hence eligibility for ICI treatments, though these classifications did not correlate with improved treatment outcomes in these groups. In contrast, TMB-high status was predictive of better ICI responses primarily in individuals of European ancestry.

The study proposed an ancestry-adjusted calibration of TMB as a means to mitigate these biases, especially in cases where matched tumor-normal sequencing data is unavailable. This recalibration demonstrated a reduction in ancestry-driven TMB inflation and yielded more accurate classifications across populations, thereby enhancing the precision of TMB as a biomarker for ICIs. While this study underscores the critical need for ancestry-diverse reference panels, the authors assessed the efficacy of their TMB recalibration approach only at the cohort level by noting the retrospective change in survival curves, without checking the effect of the recalibration in individual samples comprising the study cohorts. We identified several samples where the recalibration procedure increases the magnitude of TMB, which is inconsistent with the study’s aim of ameliorating the effect of unfiltered germline variants erroneously increasing the perceived burden to tumor mutations.

These observations motivated us to apply our previously reported method of mapping reads against pangenome reference incorporating germline variants identified in a representative cohort of healthy individuals (Rakocevic et al., 2019) to tumor sequencing samples, pursuant to the hypothesis that more accurate mapping of reads carrying germline variants will let these variants be more definitively identified, and combined with their presence in the ‘panel of normals’ used by somatic variant callers (Cibulskis et al., 2013), be filtered from the set of somatic mutations identified in the sample. We tested our approach on a small ethnically diverse cohort and present the promising results here, as motivation for future research to validate these findings in larger, ethnically diverse cohorts to refine TMB calibration techniques and ensure broader applicability in clinical settings.

## Pangenome-aware Somatic Variant Calling

We constructed a workflow (GRAF) to analyze tumor sequencing samples with the prior knowledge of known germline variants available via a relevant pangenome reference. This workflow is a variation on the standard GATK somatic calling workflow (Benjamin et al., 2019), incorporating the following changes:

i. BWA read mapper is replaced by the GRAF aligner operating with the supplied pangenome reference (passed in as a VCF file in addition to the single haplotype reference assembly used for variant calling)
ii. The standard panel of normals, comprising common germline variant repositories (dbSNP, gnomAD, ExAC, 1000 Genomes etc.), is augmented with the variants included in the pangenome. Note that the pangenome by itself is not sufficient to serve as the panel of normals because it includes only the variants characterizing the population, defined as variant appearing in a representative cohort with allele frequencies greater than a threshold (we use 5%)
iii. The somatic mutations set identified by Mutect2 variant caller is filtered by a set of static criteria designed to eliminate spurious variant calls as well as residual germline variants included in the call set.

We also constructed an analogous workflow for analyzing tumor/normal paired samples, for the purpose of generating a set of high confidence somatic mutations present in each tumor sample in our experimental dataset (Jones et al., 2015). This workflow does not require a panel of normals, instead generating the germline genotype for the patient from the normal sample.

We also created a script to calculate the tumor mutational burden (TMB) from the somatic calls, defined as the count of missense somatic mutation per mega base of genomic region (Chalmers et al., 2017).

## Evaluation of Pangenome Aware Somatic Calling Workflow

We assessed the expected accuracy of the somatic variant calls generated by GRAF workflow in two ways, by analyzing a standard benchmark sample where the somatic mutations are established with high confidence, and the by comparing the mutations identified in tumor sample against the mutations determined more accurately by analyzing the tumor normal pair in a small cohort of ethnically diverse patients.

### 1. Benchmark Accuracy

Sequencing Quality Control Phase 2 (SEQC2) study published several sequencing datasets generated from replicates of a breast cancer tissue sample, accompanied by a set of somatic mutations determined with high confidence in the tumor genome, for the purpose of benchmarking the accuracy of somatic calling methods (Fang et al., 2021). Figure 2 illustrates the overlap of SEQC2 tumor mutations in exome panel region with GRAF variant calls, plotted alongside the overlap of the GATK variant calls for comparison.

**Figure 1:**
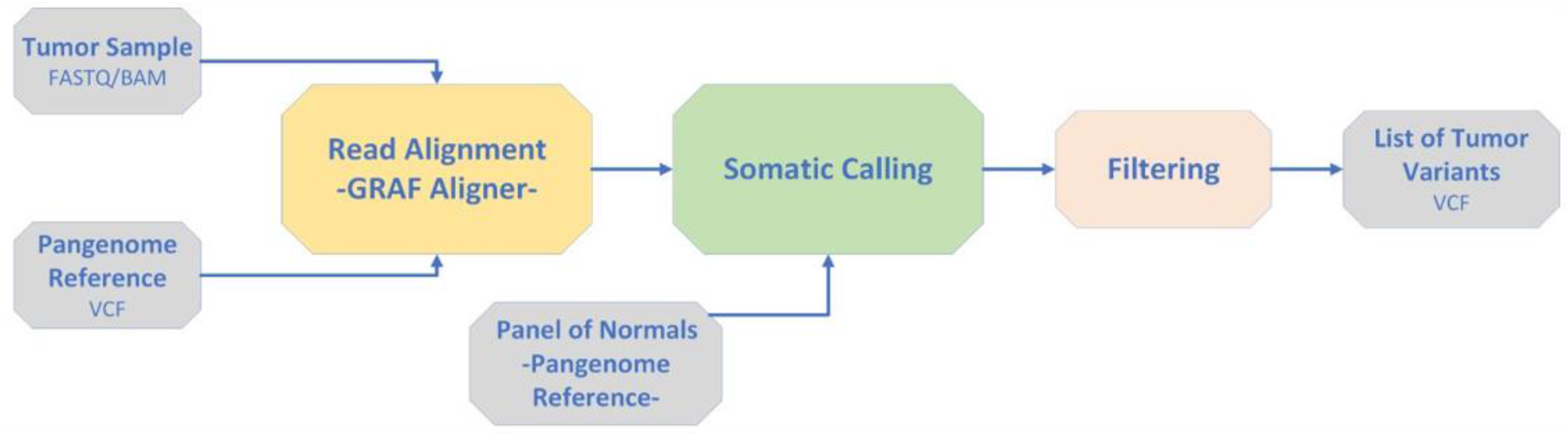
schematic of the GRAF pangenome aware somatic calling workflow

**Figure 2:**
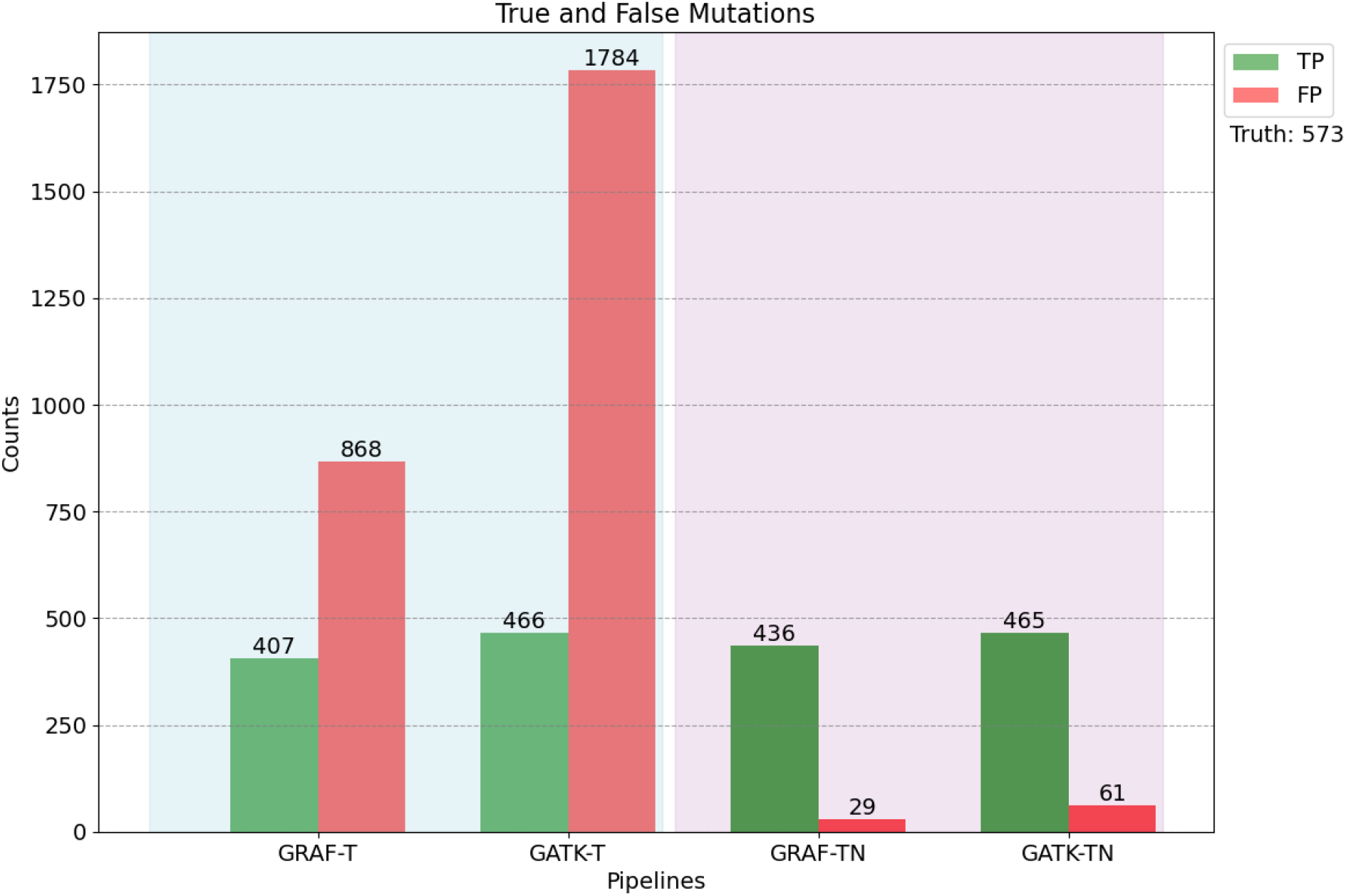
Benchmark accuracy of GRAF and GATK workflows on SEQC2 samples. Left panel shows variants called by tumor only analysis, right panel shows variants called by tumor-normal analysis

The figure compares tumor-only analysis in the left panel, demonstrating the very low precision characteristic of this method. However, GRAF results contain less than half as many false positive calls as GATK results (868 vs 1784), while showing much smaller impairment in true positive discovery (407 vs 466). The tumor-normal analysis results shown in the right panel show a drastic reduction in false positive calls while the true positive counts is close to the count from tumor only analysis, showing that

a. tumor-normal analysis is the method of choice for somatic variant calling
b. when normal sample is not available, tumor-only analysis results should be interpreted with caution due to the extremely high burden of false positive variants

This experiment also establishes that GRAF tumor-only and tumor-normal workflows are significantly more precise than the corresponding GATK workflows, at the cost of ∼10% recall impairment. However, the set of all somatic variant calls is utilized for biomarker calculations, so the loss of ∼10% extra true mutations is significantly less impactful than the gain of ∼200% spurious variant calls in tumor only analysis.

### 2. TMB estimation accuracy

Having verified that tumor-normal analysis generates more precise somatic variant calls, and that the TMB biomarker value calculated from these calls is likely to approximate the true value, we proceeded to assess the precision of TMB values estimated from 10 samples, drawn from Asian, European and African subpopulations, selected from the dataset published on TCGA by (Nassar et al., 2022). The germline sequencing data is available for all of these patients, as well as original and corrected TMB values, enabling us to carry out the tumor-normal analysis in parallel with the tumor-only analysis and compare the calculated biomarker values.

Figure 3 shows the TMB values for each sample calculated from somatic missense mutations identified by tumor-only and tumor-normal analysis, by GATK and GRAF respectively. TMB values generated by tumor-only analysis (darker bars) are similar for both GRAF and GATK workflows (mean difference 2.32, ±2.30 std), consistent with the benchmark results in experiment 1. However, TMB values by tumor-only analysis (solid bars) diverge significantly between the two workflows (mean difference 5.58, ±7.61 std), especially for samples from Asian and African subpopulations. Further, when we compare the per sample TMB values calculated by the same workflow in tumor-normal and tumor only setting, we find that the tumor only values are overestimated by both workflows, but the estimate is much more conservative in GRAF results, with mean relative increase of 35% versus 56% for GATK.

**Figure 3:**
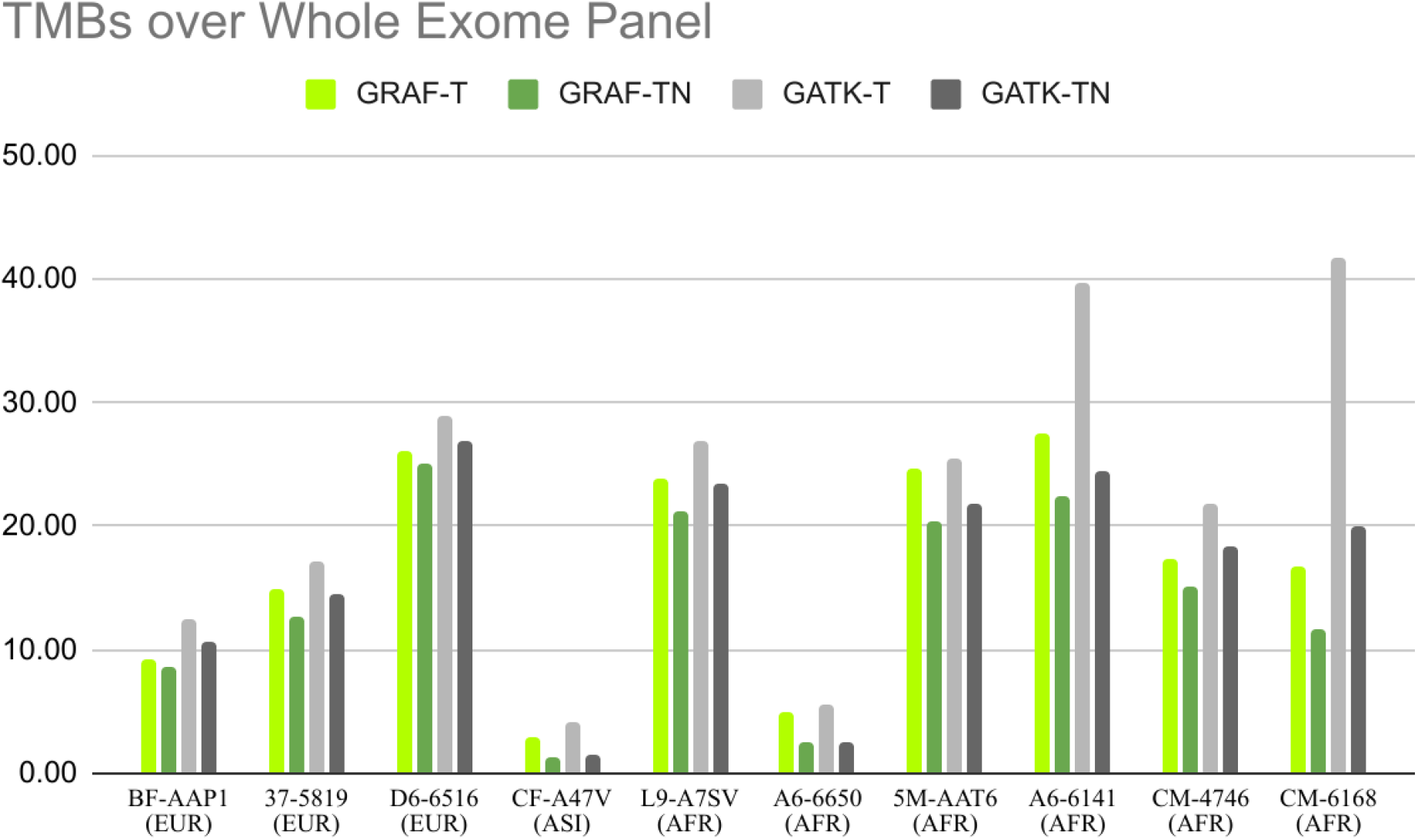
Comparison of TMB values calculated from somatic mutations identified by tumor-only analysis (solid bars) against the values calculated from tumor-normal analysis (darker bars).

These results support the hypothesis of (Nassar et al., 2022) that biomarker values are overestimated by standard somatic variant calling methods, as well as the motivating hypothesis of this study that the incorporation of population information in somatic variant calling process via pangenome references has the potential to ameliorate this bias.

### 3. Comparison against statistically recalibrated TMB values

Nasser et al. proposed a recalibration strategy to correct the population bias in calculated TMB values, by first identifying the presence of nucleotide polymorphisms in the somatic genotype as the indicator for patient’s ancestry and then applying a correction, learnt autonomously by regressing the TMB values on a large corpus of tumor normal genomes. We have the original reported TMBs as well as the corrected values available to assess the effect of the corrections and compare the values we have calculated with updated GATK and GRAF workflows. Figure 4 shows that the correction procedure leads to unexpected results for three out of 10 samples – increasing the TMB value for all European samples and setting it to zero for African samples A6-6650 and A6-6141.

**Figure 4:**
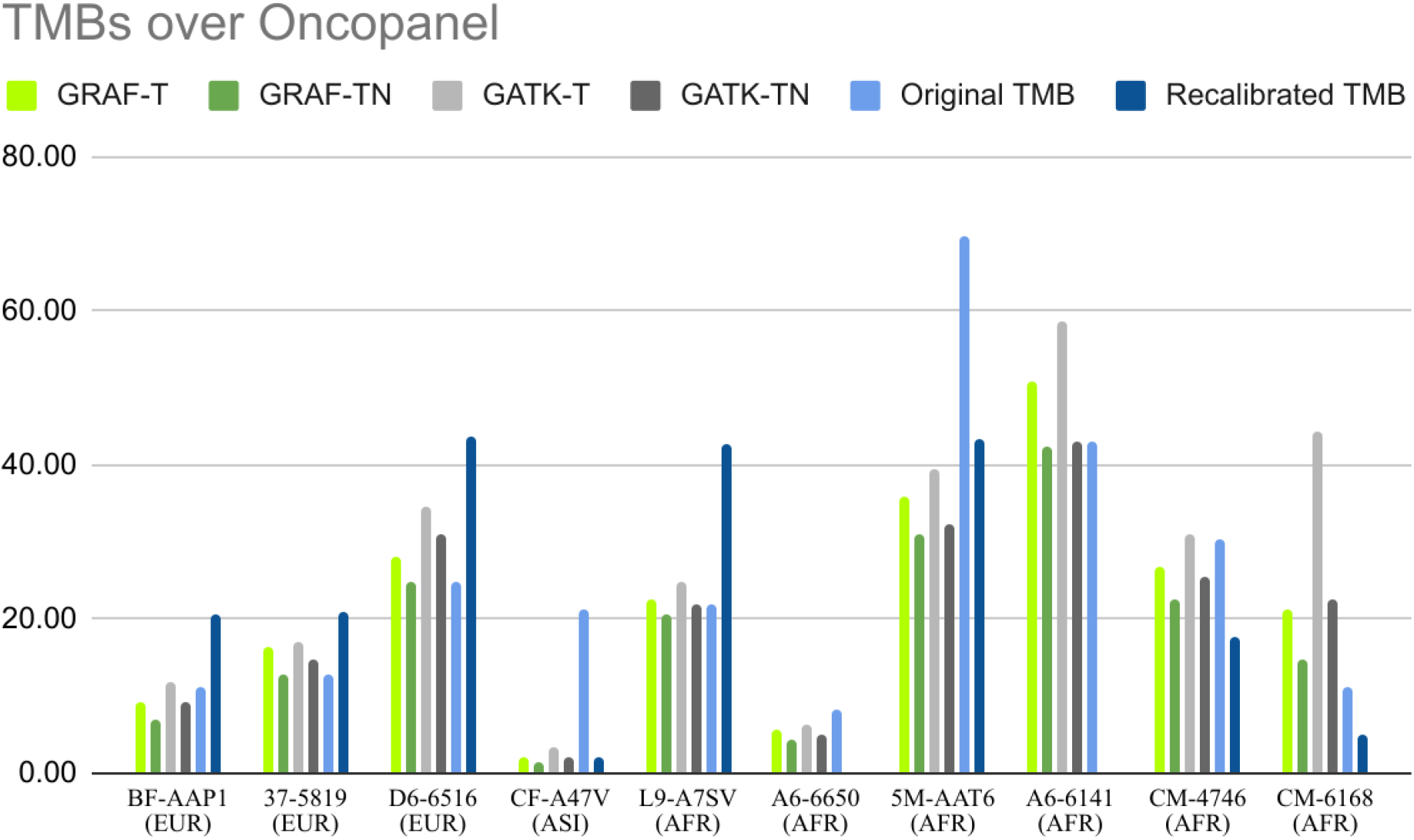
Comparison of calculated, reported and corrected TMB values over Illumina oncopanel region

### 4. Comparison of Computational Cost

Recent advances in software and hardware engineering have made pangenome based workflows competitive with germline genome sequencing analysis methods in terms of computational cost and resource burden (Rakocevic et al., 2019).

We verified that for somatic variant calling GRAF workflow is faster and cheaper than GATK workflow, costing $1.26 to process a tumor sample in 56 minutes on AWS cloud platform, versus $1.46 cost for 85 minutes. For tumor-normal analysis the time cost is higher (76 minutes costing $1.4) but also relatively more efficient than GATK (200 minutes costing $3.50).

## Conclusions and Discussion

We have presented a novel application of pangenome references to increase the precision of somatic variant calling, especially for patients belonging to non-European ancestries evolutionarily diverged from the single reference genome assembly. We assessed the accuracy of our method on a standard benchmark sample, showing significantly higher precision of tumor-only analysis, while maintaining high recall. We also showed that the computational cost of processing samples with GRAF workflow is less than the corresponding cost of GATK workflow.

We also verified the recent claims that somatic variant calls suffer from impaired precision when generated by the analysis of tumor samples harvested from patients belonging to non-European populations, leading to potential for inappropriate clinical interventions guided by biomarkers derived from the called variants. We focused on the TMB biomarker and showed that the pangenome based tumor analysis workflows have the potential to significantly reduce this bias.

As a concrete example, one tumor sample (BF-AAP1) in our test cohort resulted in TMB value of ∼12 when analyzed with the standard GATK workflow, but pangenome-based GRAF workflow reduced the count of missense mutations leading to calculated TMB getting reduced to 9, which is closer to the value calculated by the more precise tumor-normal analysis of the same sample. This difference is clinically significant as TMB=10 is often used as the threshold for decision to administer ICI therapy.

The value of pangenome references has been established for improved understanding of germline genomic variation (Liao et al., 2023), (Tetikol et al., 2022). We hope that our results, demonstrating clinically relevant accuracy available at a lower processing cost, will encourage further studies exploring the value of pangenome references for better understanding of cancer genomes and eventually lead to their widespread adoption for processing of cancer sequencing datasets.

GRAF somatic calling workflow is available on the Cavatica computational platform (Berke et al., 2024) maintained by Velsera Inc.

## Acknowledgements

Brandi Davis-Dusenbery provided the initial hypothesis for this study.

Renee L Sears contributed insights for effective filtering of spurious mutations.

Authors appreciate <u>Michael Hultner’s</u> support and enablement for this study, and <u>Jack DiGiovanna’s</u> continued evangelism of the promise of pangenomes including this somatic application.

## Appendix

### Data for the figures

*TMB values calculated over WES region (Figure 3)*

| Samples | GRAF |  | GATK |  |
| --- | --- | --- | --- | --- |
|  | T | TN | T | TN |
| TCGA-BF-AAP1 | 9.33 | 8.61 | 12.57 | 10.74 |
| TCGA-37-5819 | 14.98 | 12.70 | 17.19 | 14.52 |
| TCGA-D6-6516 | 26.06 | 25.03 | 29.02 | 26.96 |
| TCGA-CF-A47V | 2.93 | 1.39 | 4.11 | 1.49 |
| TCGA-L9-A7SV | 23.82 | 21.25 | 26.96 | 23.44 |
| TCGA-A6-6650 | 5.04 | 2.60 | 5.47 | 2.54 |
| TCGA-5M-AAT6 | 24.65 | 20.43 | 25.49 | 21.92 |
| TCGA-A6-6141 | 27.45 | 22.44 | 39.65 | 24.41 |
| TCGA-CM-4746 | 17.27 | 15.16 | 21.92 | 18.38 |
| TCGA-CM-6168 | 16.76 | 11.62 | 41.79 | 19.92 |

*TMB values calculated over oncopanel region (Figure 4)*

| Samples | GRAF |  | GATK |  | Nasser et al |  |
| --- | --- | --- | --- | --- | --- | --- |
|  | T | TN | T | TN | Original | Recalibrated |
| TCGA-BF-AAP1 | 9.18 | 7.06 | 12.00 | 9.18 | 11.35 | 20.73 |
| TCGA-37-5819 | 16.24 | 12.71 | 16.94 | 14.82 | 12.86 | 20.94 |
| TCGA-D6-6516 | 28.24 | 24.71 | 34.59 | 31.06 | 24.96 | 43.68 |
| TCGA-CF-A47V | 2.12 | 1.41 | 3.53 | 2.12 | 21.18 | 1.99 |
| TCGA-L9-A7SV | 22.59 | 20.47 | 24.71 | 21.88 | 21.94 | 42.73 |
| TCGA-A6-6650 | 5.65 | 4.24 | 6.35 | 4.94 | 8.32 | 0.00 |
| TCGA-5M-AAT6 | 36.00 | 31.06 | 39.53 | 32.47 | 69.59 | 43.25 |
| TCGA-A6-6141 | 50.82 | 42.35 | 58.59 | 43.06 | 43.12 | 0.00 |
| TCGA-CM-4746 | 26.82 | 22.59 | 31.06 | 25.41 | 30.26 | 17.81 |
| TCGA-CM-6168 | 21.18 | 14.82 | 44.47 | 22.59 | 11.35 | 5.08 |

